# Drug Conjugates of Antagonistic RSPO4 Mutant For Simultaneous Targeting of LGR4/5/6 for Cancer Treatment

**DOI:** 10.1101/2021.02.05.429956

**Authors:** Jie Cui, Soohyun Park, Wangsheng Yu, Kendra Carmon, Qingyun J. Liu

## Abstract

LGR4-6 (Leucine-rich repeating containing, G-protein-coupled receptors 4, 5, and 6) are three related receptors with distinct roles in organ development and stem cell survival. All three receptors are upregulated in gastrointestinal cancers to different levels, and LGR5 has been shown to be enriched in cancer stem cells. Antibody-drug conjugates (ADCs) targeting LGR5 showed robust antitumor effect in vivo but could not eradicate tumors due to plasticity of LGR5-positive cancer cells. As LGR5-negative cells often express LGR4 or LGR6 or both, we reasoned that simultaneous targeting of all three LGRs may provide a more effective approach. R-spondins (RSPOs) bind to LGR4-6 with high affinity and potentiate Wnt signaling. We identified an RSPO4 mutant (Q65R) that retains potent LGR binding but no longer potentiates Wnt signaling. The RSPO4 mutant was fused to the N-terminus of human IgG1-Fc to create a peptibody which was then conjugated with cytotoxins monomethyl auristatin or duocarmycin by site-specific conjugation. The resulting peptibody drug conjugates (PDCs) showed potent cytotoxic effects on cancer cell lines expressing any LGR in vitro and suppressed tumor growth in vivo without inducing intestinal enlargement or other adverse effects. These results suggest that RSPO-derived PDCs may provide a novel approach to the treatment of cancers with high LGR expression.

## Introduction

LGR4-6 (leucine-rich repeat containing, G protein-coupled receptor 4, 5, and 6) are three related membrane receptors with a large extracellular domain (ECD) and a seven transmembrane (7TM) domain typical of GPCRs (1,2). In normal adult tissues, LGR4 is expressed broadly at low to moderate levels whereas LGR5 and LGR6 are mostly restricted to adult stem cells in the gastrointestinal system and skin (3–6). The three receptors are often co-upregulated in various cancer types, particularly in cancers of the gastrointestinal system, with LGR4 co-expressed with LGR5 or LGR6 (7–12). Furthermore, LGR5 has been shown to be enriched in cancer stem cells (13–16) and LGR5-positive cancer cells were shown to fuel the growth of tumor mass and metastasis (17,18). LGR6 is a marker of cancer stem cells of squamous carcinoma (16). TCGA’s RNA-Seq data of colon, liver, and stomach cancers confirmed that LGR4 expression at high levels in nearly all cases while LGR5 and LGR6 were co-expressed with LGR4 in the majority of colon cancer and substantial fractions of liver and stomach cancer. LGR5 is also a marker of stem cells in the liver and highly expressed in hepatocellular carcinomas (HCC) with β-catenin mutations (19,20).

R-spondins are a group of four related secreted proteins (RSPO1-4) that play critical roles in normal development and survial of adult stem cells (21). RSPOs function as ligands of LGR4-6 to potentiate Wnt signaling and support stem cell growth (22–25). Mechanistically, RSPO-LGR4 form a complex to inhibit the function of RNF43 and ZNRF3, two E3 ligases that ubiquitinate Wnt receptors for degradation, leading to higher Wnt receptor levels and stronger signaling (26,27). In constrast, RSPO-LGR5 potentiates Wnt signaling through a different mechanism and although RSPO4 has been shown to bind LGR5 it is unable to activate LGR5-mediated Wnt signaling (28). All four RSPOs comprise a N-terminal furin-like domain with 2 repeats (Fu1 and Fu2) and thrombospondin-like domain (TSP) at the C-terminus (21). The furin-like domain is both necessary and sufficient to bind LGRs to potentiate Wnt signaling whereas the TSP domain enhances the potency of furin-like domain (29–31). Crystal structure analysis revealed that the Fu1 domain binds to the E3 ligases while the Fu2 domain binds to the extracellular domain of LGR (32–35).

Antibody–drug conjugates (ADCs) are monoclonal antibodies (mAbs) that are covalently linked to cell-killing cytotoxins (payloads). This approach combines the high specificity of mAbs with potent cytotoxic drugs, creating “armed” antibodies that can directly deliver the payload to antigen-enriched tumor cells while minimizing systemic toxicity (36–38). We and others have shown that anti-LGR5 ADCs are highly effective in inhibiting the growth of LGR5-positive colon cancers in xenograft without major adverse effect (12,39). However, tumors eventully came back due to plasticity of colon cancer cells (12,18,39), which have been shown to interconvert between an LGR5-positive and LGR5-negtive phenotype (14). Both LGR5-positive and LGR5-negative colon cancer cells have been shown to express LGR4 (40). Given that RSPOs bind to all three LGRs with high affinity, we reasoned that drug conjugates of an RSPO mutant protein that maintains binding to LGR without activating Wnt signaling would be able to target cancer cells expressing any LGR and therefore overcome plasticity of cancer cells. Here we report that an RSPO4-furin domain mutant Gln65 to Arg (Q65R) fused to human IgG1-Fc (peptibody) and conjugated with different cytotoxins was able to inhibit the proliferation of LGR-positive cancer cells in vitro and tumor growth in vivo. Importantly, this RSPO mutant peptibody drug conjugate (PDC) was well tolerated in vivo and it did not induce hyperproliferation of crypt stem cells, a hallmark of RSPO agonism in the intestine (41).

## Results and Discussion

### Characterization of RSPO4-Fu mutant with high-affinity binding to LGR4-6 and antagonist activity in Wnt signaling

Since RSPO4 has the highest binding affinity for LGR4-6 yet the lowest functional activity in potentiating Wnt signaling (22,42,43), we reasoned that RSPO4-Fu may be the best starting ligand for development of a PDC. Based on co-crystal structures of RSPO1/2 furin domain with RNF43/ZNRF3-ECD (Extracellular Domain) and LGR5-ECD (32,33), Gln65 (Q65) of RSPO4 is predicted to interact with RNF43/ZNRF3 directly (Fig. 1A). Homozygous mutation of RSPO4 Q65R in human causes nail formation defects (44,45), and introduction of this mutation to the corresponding Gln residues in RSPO1 (Q71) and RSPO2 (Q75) led to near total loss of activity in potentiating Wnt signaling (34,43). We expressed and purified RSPO4-Fc fusion proteins comprising RSPO4-Fu domain (R4Fu, AA27-124) of either wild-type (WT) or Q65R fused to the N-terminus of human IgG1-Fc. The two proteins, designated R4Fu-WT and R4Fu-Q65R, were purified and showed similar high affinity binding to LGR4- and LGR5-overexppressing HEK293T cells (Fig. 1B). Importantly, the mutant was completely inactive in potentiating Wnt/β-catenin signaling whereas the WT protein showed robust, dose-dependent activity in STF cells which express LGR4 endogenously (Fig. 1C). Since RSPO1 and RSPO2 have complete and partial dependence on LGR4 for potentiating Wnt signaling, respectively (31,46), R4Fu-Q65R is expected to be able to bind to LGR4 and prevent the binding of RSPO1, and therefore antagonize RSPO1/2 activity. Indeed, R4Fu-Q65R antagonized both RSPO1 and RSPO2 activity with better potency and efficacy on RSPO1 (Fig. 1D).

**Figure 1.**
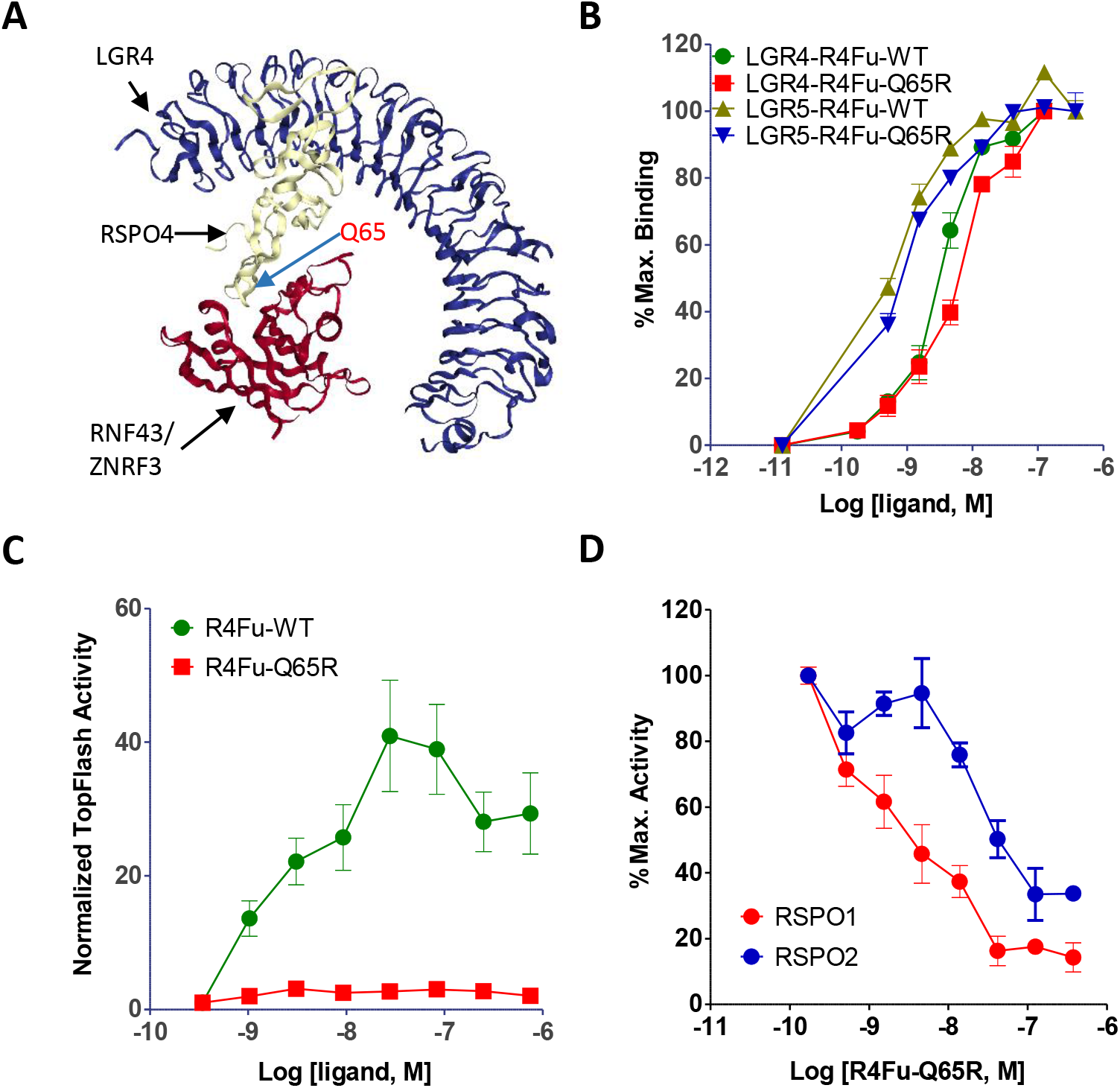
RSPO4-Fu-Q65R retained high affinity binding to LGR4/5 but lost Wnt-potentiating activity. A, A proposed structure of RSPO4Fu binding to RNF43/ZNRF3 and LGR4 modeled after LGR5ECD-RSPO2Fu-ZNRF3 (PDB 4UFS). B, Binding analysis of purified R4Fu-WT and - Q65R to HEK293 cells overexpressing LGR4 or LGR5. C, TOPFlash Wnt/β-catenin signaling activity of R4Fu-WT and - Q65R in HEK293-STF cells. D, Antagonist activity of R4Fu-Q65R on RSPO1 and RSPO2 in TOPFlash Wnt/β-catenin signaling assay.

### R4Fu-Q65R conjugated with MMAE or duocarmycin showed potent and specific effect on LGR-high cancer cell lines

To evaluate the potential of the R4Fu-Q65R peptibody as a drug carrier, site-specific conjugation was performed with a cleavable linker-cytotoxin as schematically illustrated (Fig. 2A). In brief, R4Fu-Q65R incorporating an LLQGA tag at the C-terminus was expressed and purified with the Q (Gln) residue in the LLQGA tag serving as the recipient group for transgluminase (47). PEG4-VC-PAB-MMAE (ethylene glycol-4x-valine-citruline-p-aminobenzoyloxycarbonyl-MMAE) was incubated with the peptibody in the presence of bacterial transglutaminase as described previously (48). The PDC product, designated R4Fu-Q65R-MMAE, was purified and confirmed by gel analysis (Fig. 2B). Mass spectrometry analysis performed validated the conjugation was specific to the LLQGA tag (Fig. 2C). A similar PDC using duocarmycin SA (DMSA) as payload was also generated, and the product was designated R4Fu-Q65R-DMSA.

**Figure 2.**
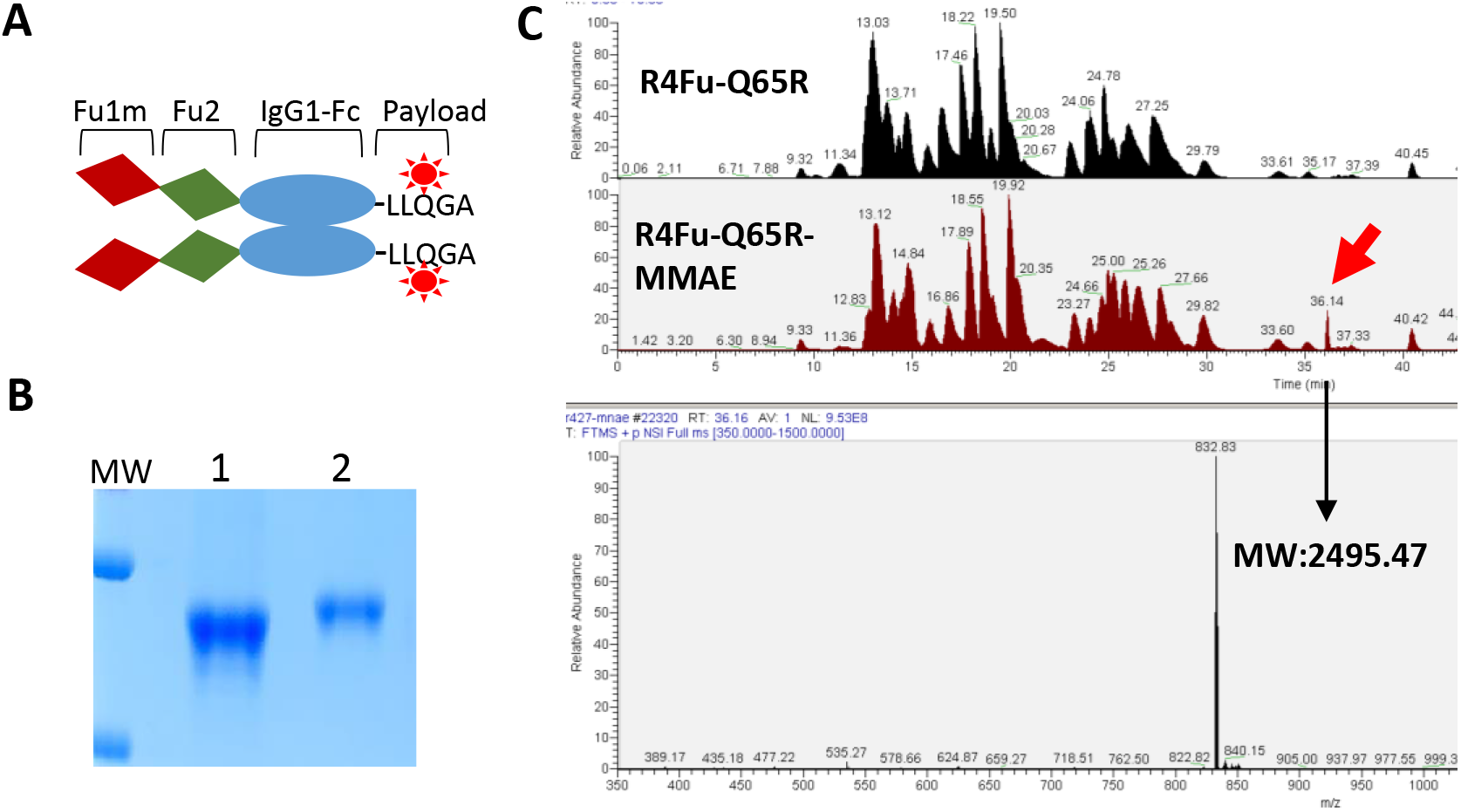
Site-specific conjugation of MMAE to R4Fu-Q65R. A, a schematic diagram of the R4Fu-Q65R with an LLQGA tag at the C-terminus. B., Coomassie blue staining of R4Fu-Q65R before (lane 1) and after conjugation (lane 2). C, LC-MS analysis of R4Fu-Q65R before and after conjugation.

We first tested cytotoxic activity of R4Fu-Q65R-MMAE on HEK293T cells overexpressing LGR4, LGR5, or LGR6. The PDC showed much more potent cytotoxic effect on HEK293T overexpressing the LGRs when compared to parental HEK293T cells (Fig. 3A), which express LGR4 endogenously at levels sufficient for enhancing Wnt signaling. When tested on the LoVo colon cancer cell line, which expresses both LGR4 and LGR5, R4Fu-Q65R-MMAE inhibited cell growth with an IC50 of ~1 nM. Knockout (KO) of LGR4 (which suppressed expression of LGR5) or knockdown (KD) of LGR5 led to loss of sensitivity to the PDC (Fig. 3B), indicating the cytotoxicity was mediated by LGR4 and LGR5. We also compared the R4Fu-Q65R PDCs with anti-LGR5 ADC (39) and anti-LGR4 ADC side-by-side on LoVo cells. Both PDCs were ~3x more potent than the anti-LGR5 ADC and ~10x more potent than the anti-LGR4 ADC (Fig. 3C). The higher potencies of PDCs, despite similar affinity of PDCs and ADCs in binding to the receptors, may be due to simultaneous targeting of two receptors and/or differential intracellular trafficking induced by ligand versus antibody. We then evaluated the two PDCs against a large panel of LGR-high cancer cell lines of the digestive system, including DLD1 and HT29 (colon), HepG2 and PLC-PRF5 (liver cancer), and AGS (stomach) and obtained IC50s between 1-10 nM, with data shown for PLC and AGS shown in Figure 3D. These results indicate that RSPO4-Fu PDCs are able to inhibit the growth of a wide range of cancer cell lines expressing any LGR at sufficient levels.

**Figure 3.**
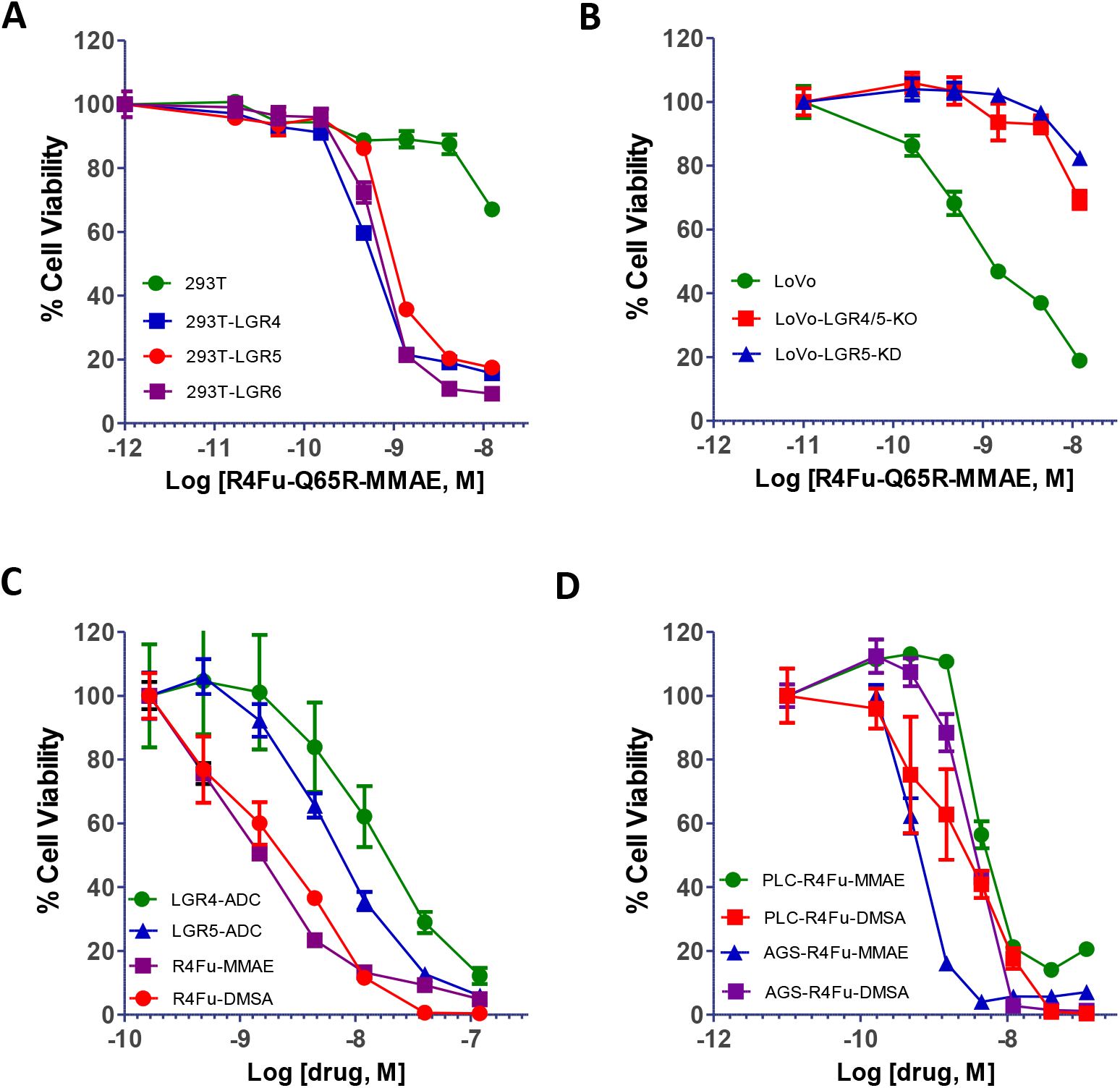
Cytotoxic activity of R4Fu-Q65R-MMAE and –DMSA in various cell lines. A, Viability of HEK293T cells with or without over-expressing LGR4, LGR5, or LGR6. B, Viability of LoVo cells with KO of LGR4/5 or KD of LGR5 treated with R4Fu-Q65R PDC. C, Viability of LoVo cells treated with ADCs of LGR4, LGR5, or R4Fu-Q65R PDC. D, Viability of PLC/PRF5 and AGS cells treated with R4Fu-Q65R PDC.

### Anti-tumor activity of R4Fu PDC in vivo

We tested R4Fu-Q65R-MMAE and R4Fu-65R-DMSA in a xenograft model of LoVo cells in athymic nude mice. When tumors reached the size of ~130 mm3, the animals were randomized into 4 groups with 5-6 per group. The animals were administered PBS (vehicle), unconjugated R4Fu-Q65R, R4Fu-Q65R-MMAE, or R4Fu-Q65R-DMSA at 5 mg/kg by intraperitoneal injection every other day for a total of eight injections. Tumor sizes were measured and animals were monitored for general health. Both MMAE and DMSA - conjugated PDC showed significant anti-tumor effect with 70-80% inhibition of tumor growth (Fig. 4A) and significant increase in survival (Fig. 4B, survival data not shown for unconjugated for clarity). Neither body weight loss nor other gross toxicity was observed with any of the treatment groups.

**Figure 4.**
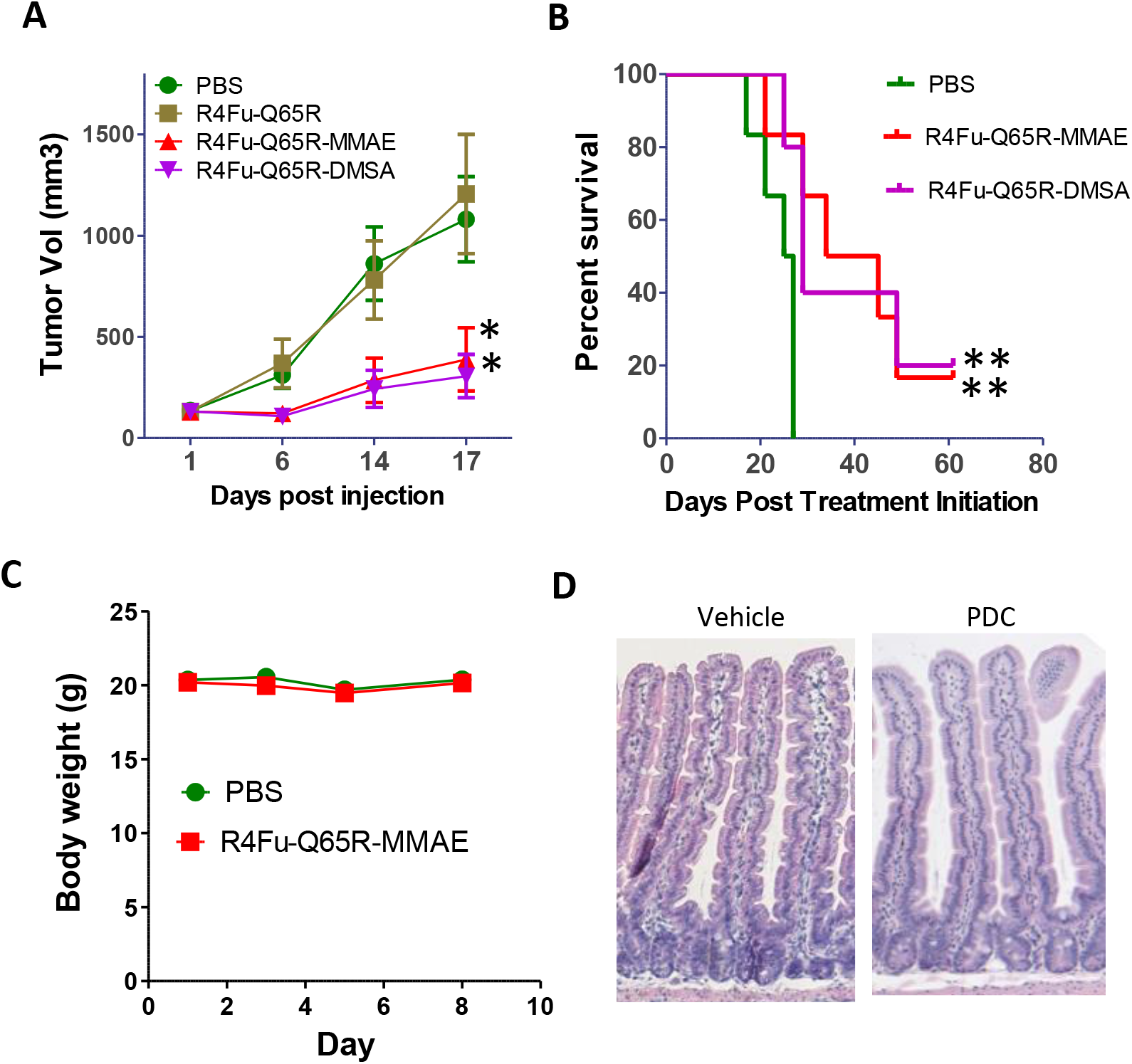
Anti-tumor effect of R4Fu-Q65R PDCs in xenograft models of LoVo cells and in vivo toxicity. A, Tumor growth curve. *p< 0.05 vs PBS group. B, Kaplan-Meir survival plot of R4Fu-Q65R-MMAE and –DMSA vs PBS. **p= 0.02 vs PBS, Log-rank test. Last treatment was on Day 17. C, Body weights of C57/Bl mice dosed with vehicle ((PBS) or PDC at 15 mg/kg on Days 1, 3, and 5 (N =3 per group). D, H&E staining of small intestine at termination (Day 8) from vehicle or PDC-treated animals.

### R4Fu-Q65R-MMAE was well tolerated in vivo

To evaluate tolerability of the PDCs, R4Fu-Q65R-MMAE was given to normal C57/Bl mice for one week and monitored body weight and general well-being. At up to 15mg/kg, the animals showed no obvious signs of toxicity or significant loss of body weight (Fig. 4C). Blood analysis at termination showed no significant change in blood counts and liver function enzymes. H&E staining of intestines revealed no effect on crypt size and epithelium histology (Fig. 4D). Importantly, the lack of effect on crypt size indicates that the PDC at this dose level was not able to antagonize the activity RSPO2 and RSPO3 which are essential for self-renewal and proliferation of crypt stem cells (49), most likely because RSPO2/3 could partially function without LGR (31,46). It also suggests that the combined LGR levels in intestinal stem cells are too low to mediate sufficient cytotoxicity delivered by the PDCs. These data indicate that R4Fu PDCs are well-tolerated at 3 times the therapeutically effective dose.

In summary, we have created a RSPO4-Fu mutant that is defective in potentiating Wnt signaling but retains high affinity binding to LGRs. An Fc fusion protein of this mutant was able to antagonize RSPO function and deliver cytotoxic drugs into LGR-high cancer cells to achieve robust anti-tumor effect in vitro and in vivo. Furthermore, the PDC displayed more potent cytotoxic activity than ADCs targeting LGR4 or LGR5 alone. Preliminary toxicology studies also revealed that the PDC was well-tolerated at 3 times the efficacious dose. These data provided proof-of-concept for the use of a functionally-inactive ligand peptibody for competitive receptor antagonism and drug delivery. The approach offers the potential of targeting all three LGRs simultaneously and may provide a breakthrough concept to the treatment of a significant proportion of cancers of the gastrointestinal system that co-express high levels of LGR4-6. The PDCs still need improvement in efficacy and potency as well as complete characterization in pharmacokinetics and safety.

## Materials and Methods

### Cell lines and plasmids

LoVo cells were obtained from the laboratory of Dr. Shao-Cong Sun at MD Anderson Cancer Center (Houston, TX). Knockout of LGR4 in LoVo cells were carried out uisng the CRISPR/Cas9 lentiviral vector as described (31,50). All other cancer cell lines were purchased from ATCC (Americal Tissue Culture Collection) and cultured as specified. HEK293 cells expressing LGR4 or LGR5 or LGR6 were generated as previously reported (22,25).

### Protein expression, purification, and characterizaiton

R4Fu-WT and R4Fu-Q65R were cloned into the vector pCEP5 using standard cloning methods. Protein production and characterizaiton were carried out as we decribed previously (31). Receptor binding, Wnt signalign assays (TOPFlash) were also performed as before (31).

### ADC synthesis and characterization

NH2-PEG4-VC-PAB-MMAE was purchased from Levena Biopharma (San Diego, California). Conjugation to R4Fu-Q65R was carried out using bacterial transgluminase as described by Strop et al (48).

### In vitro cell viability assays

Cells were seeded at various numbers (depending on growth rate) in 96-well plates and serial dilutions of PDC or ADCs were added. The cells were incubated for 4 days and cell viabilty was measured using CellTiter-Glo® Luminescent Cell Viability Assay (Promega).

### In vivo xenograft studies

Animal studies were carried out in strict accordance with the recommendations of the Institutional Animal Care and Use Committee of the respetive institutes. For LoVo xenograft study, female 9-week-old nu/nu mice (Charles River Laboratories) were subcutaneously inoculated with 5 × 106 cells in 1:1 mixture with Matrigel (BD Biosciences). After 3 weeks, when tumor size reached approximately ~100 mm3, mice were randomized into 5 groups of 6 mice per group and given vehicle (PBD), unconjuated R4Fu-Q65R, R4Fu-Q65R-MMAE or –DMSA once every other day at 5 mg/kg by intraperitoneal injection for a total of 8 doses. For tolerability study in normal animals, 11-week old female C57BL/6 mice were given a single dose of PBS or R4Fu-Q65R-MMAE at 15 mg/kg by IV injection. The animals were monitored for body weight and general behavior for 10 days. Clinical chemistry was carried out by Veterinary Laboratory Medicine facility at the Department of Veterinary Medicine & Surgery, MD Anderson Cancer Center. Intestines were harvested and examed following H&E staining.

### Statistical analysis

Data are expressed as mean ± SEM or SD as indicated in the Results section. Multiple comparisons used one-way ANOVA and Dunnett’s post hoc analysis. P ≤ 0.05 was considered statistically significant.

## Ackledgement

The authors would like to thank Ms. Li Li and Dr. Sheng Pan of the Clinical and Translational Proteomics Service Center at UTHealth for the mass spectrometry analysis and Dr. Shao-cong Sun of M.D. Anderson Cancer Center for the LoVo cells. This project was supported in part by funding from Wntrix Inc (to QJL) and the Janice David Gordon for Bowel Cancer Research Endowment (to QJL).

